# Growth and division mode plasticity is dependent on cell density in marine-derived black yeasts

**DOI:** 10.1101/2021.05.16.444389

**Authors:** Gohta Goshima

## Abstract

The diversity and ecological contribution of the fungus kingdom in the marine environment remain under-studied. A recent survey in the Atlantic (Woods Hole, MA, USA) brought to light the diversity and unique biological features of marine fungi. The study revealed that black yeast species undergo an unconventional cell division cycle, which has not been documented in conventional model yeast species such as *Saccharomyces cerevisiae* (budding yeast) and *Schizosaccharomyces pombe* (fission yeast). The prevalence of this unusual property is unknown. Here, I collected and identified 65 marine fungi species across 40 genera from the surface ocean water, sediment, and the surface of macroalgae (seaweeds) in the Pacific (Sugashima, Toba, Japan). The Sugashima collection largely did not overlap with the Woods Hole collection and included several unidentifiable species, further illustrating the diversity of marine fungi. Three black yeast species were isolated, two of which were commonly found in Woods Hole (*Aureobasidium pullulans, Hortaea werneckii*). Surprisingly, their cell division mode was dependent on cell density, and the previously reported unconventional division mode was reproduced only at a certain cell density. For all three black yeast species, cells underwent filamentous growth with septations at low cell density and immediately formed buds at high cell density. At intermediate cell density, two black yeasts (*H. werneckii* and an unidentifiable species) showed rod cells undergoing septation at the cell equator. In contrast, all eight budding yeast species showed a consistent division pattern regardless of cell density. This study suggests the plastic nature of the growth/division mode of marine-derived black yeast.

## Introduction

Understanding the habitats of marine organisms is important for understanding marine ecology. Much of this effort has been put on macro-organisms, such as fishes and benthic invertebrates, as well as unicellular microorganisms such as algae and bacteria (OBIS, https://obis.org/). Relatively little attention has been paid to fungi, and little is known about their life cycle and physiology (Amend et al., 2019; Gladfelter et al., 2019). The genetics and cell biology of these organisms have been pioneered in a few terrestrial fungal systems, such as *Saccharomyces cerevisiae, Schizosaccharomyces pombe*, and *Aspergillus nidulans* (Feyder et al., 2015; Nurse and Hayles, 2019; Osmani and Mirabito, 2004). An important step towards understanding the ecology of marine fungi is the identification of fungi in various locations.

An interesting study was published in 2019, in which 35 fungi were collected from the sediment, surface ocean water, and benthic animals (corals, sponges) around Woods Hole, MA, USA (Mitchison-Field et al., 2019). That study investigated the division pattern of several black yeast species, which is a polyphyletic group of fungi that has melanised cell walls. Black yeasts have been of ecological interest, as they are tolerant to extreme environmental conditions such as super-high salinity (Gostincar et al., 2012; Moreno et al., 2018). Live imaging of black yeasts uncovered remarkable unconventional features in their cell division cycle (Mitchison-Field et al., 2019). For example, a single *Hortaea werneckii* cell always underwent fission during its first cell division, but the next division involved budding at 92% probability. This observation challenged the conventional view that a single cell division pattern is intrinsic to a yeast species; for example, *S. pombe* and *S. cerevisiae* always divide via fission or budding, respectively. In another striking example, more than 50% of *Aureobasidium pullulans* cells produced multiple rounds of simultaneous buds, which is never observed in *S. cerevisiae* (Mitchison-Field et al., 2019). From an ecological point of view, this study urges the necessity of further sampling and characterisation, as what has been reported to date is unlikely to be the full set of marine fungi in the ocean.

In this study, inspired by (Mitchison-Field et al., 2019), free-living marine fungi were collected in front of Nagoya University’s Marine Biological Laboratory (NU-MBL) on Sugashima Island, Toba, Japan (Fig. 1A). The species were identified via DNA barcode sequencing, and the division pattern was observed for budding and black yeasts (no fission yeast was isolated). The collected species, or even genera, only partially overlapped with those identified by (Mitchison-Field et al., 2019), suggesting the existence of highly divergent fungal species in the ocean. Surprisingly, the division pattern of black yeasts (*H. werneckii, A. pullulans*, and other unidentified species) was initially inconsistent with those previously reported in (Mitchison-Field et al., 2019), and this enigma was solved through the observation of cell density-dependent alterations in their division patterns.

**Figure 1.**
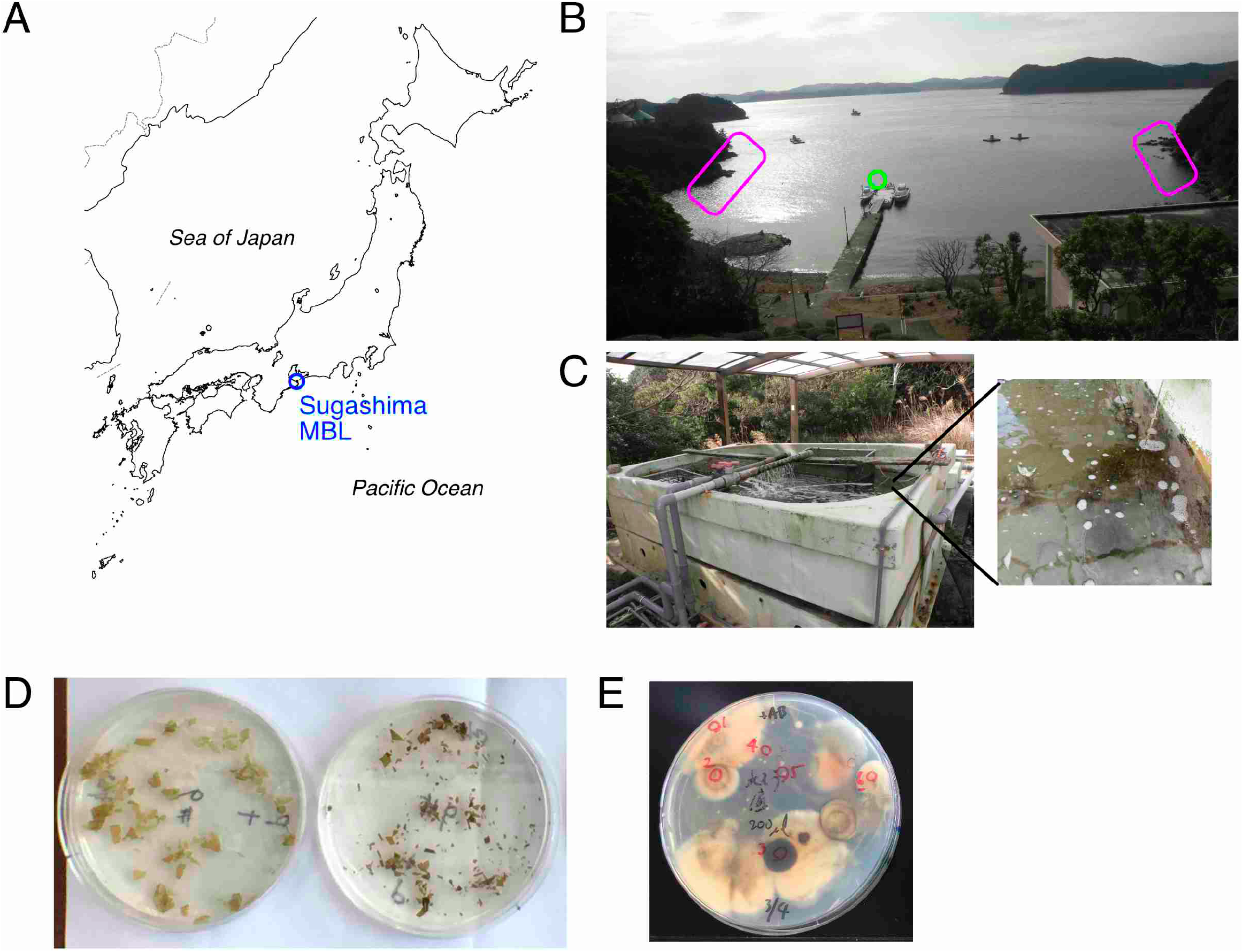
Collection of marine fungi from surface ocean water, sediment, and seaweeds. (A) Location of Sugashima Marine Biological Laboratory (NU-MBL). (B) Surface ocean water was obtained at the pier (green), whereas seaweeds were collected at the intertidal zone (magenta). (C) Outdoor tank at NU-MBL which has a continuous flow of unfiltered sea water. The sediment and surface water were the sources of marine fungi. (D) Severed seaweeds on the fungal medium plate. Several fungal colonies grew after several days. (E) Examples of fungal colonies on the plate (sediment sample). Each colony was marked and subjected to genotyping PCR and transfer to a fresh plate.

## Results

### Isolation of fungi at the Sugashima marine laboratory

Fungi were isolated from four sources: surface ocean sea water near the beach (Fig. 1B, green), an outdoor tank that has various benthic animals and seaweeds (circulated surface water and sediment; Fig. 1C), and sliced seaweeds (each ∼3 cm) that were collected in the intertidal zone in front of the laboratory (Fig. 1B [magenta], D). The sediment was most enriched with fungi (Fig. 1E; 200 µL sediment). Fungal colonies were obtained from 100 mL surface sea water, and 7.5 ± 8.5 fungal colonies were obtained (n = 6), whereas seaweeds were a more abundant source of fungi (25 ± 26 colonies isolated from a ∼3 cm piece of seaweed body [n = 10]). However, based on colony colour and shape, it was deduced that many colonies were derived from the same species. In total, 74 colonies were regrown on separate plates (15 examples are shown in Fig. 2), and two (ITS1/4 and NL1/4) or more barcode regions were sequenced (Table 1, Table S1). For some clones, the species or even genera could not be identified because of the high deviation of the barcode sequences from the known sequences registered in the database (named with “sp.” or described as “unclear” in Table 2). The total number of species collected was 65 (Table 2).

**Figure 2.**
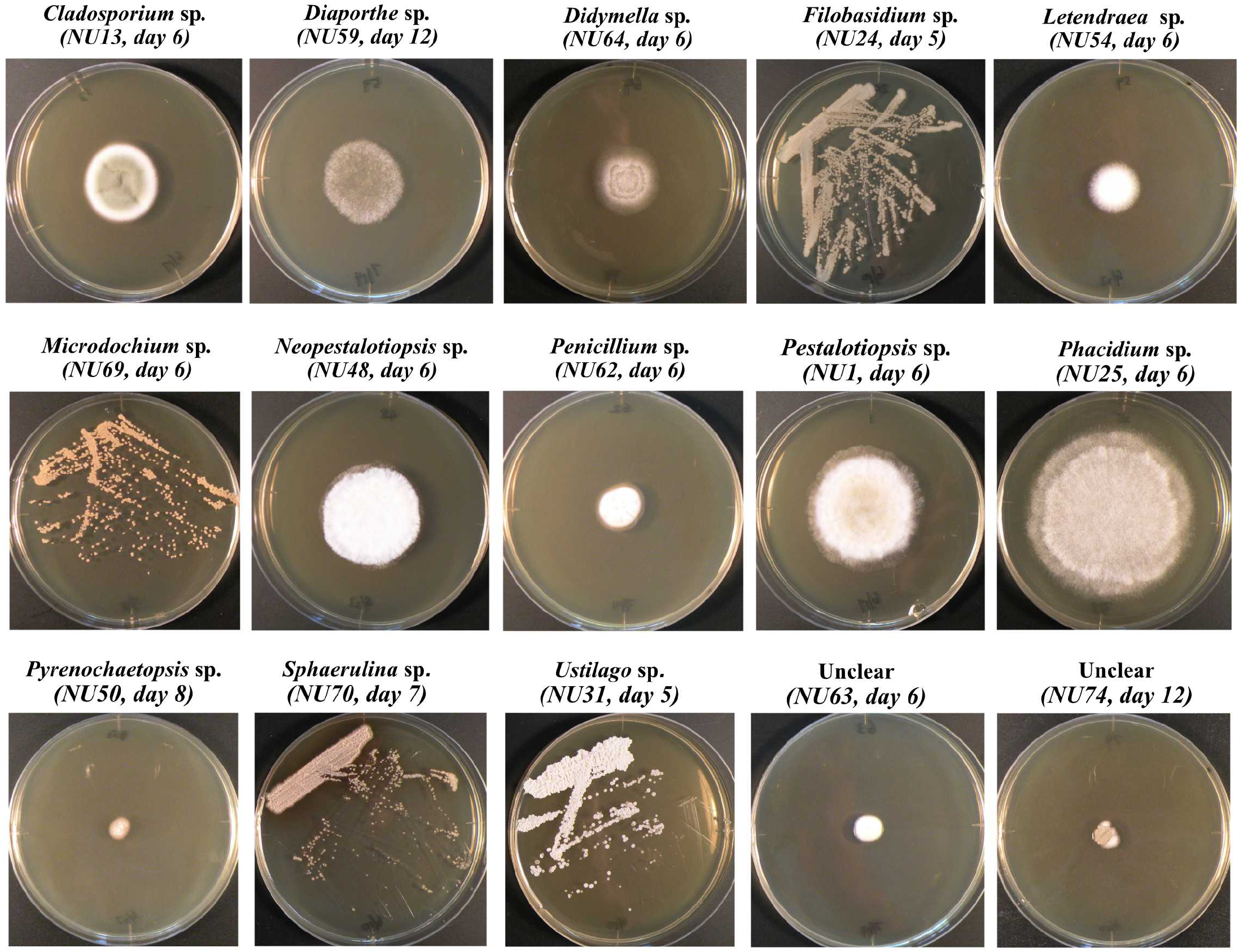
Examples of fungal colonies. Fifteen fungal colonies, whose identity could not be determined, are shown. Filamentous fungi were inoculated onto the centre of the YPD plate, whereas yeasts were streaked. They were incubated at 18°C for indicated days. The diameter of the plates in this figure is 9 cm.

**Table 1:**
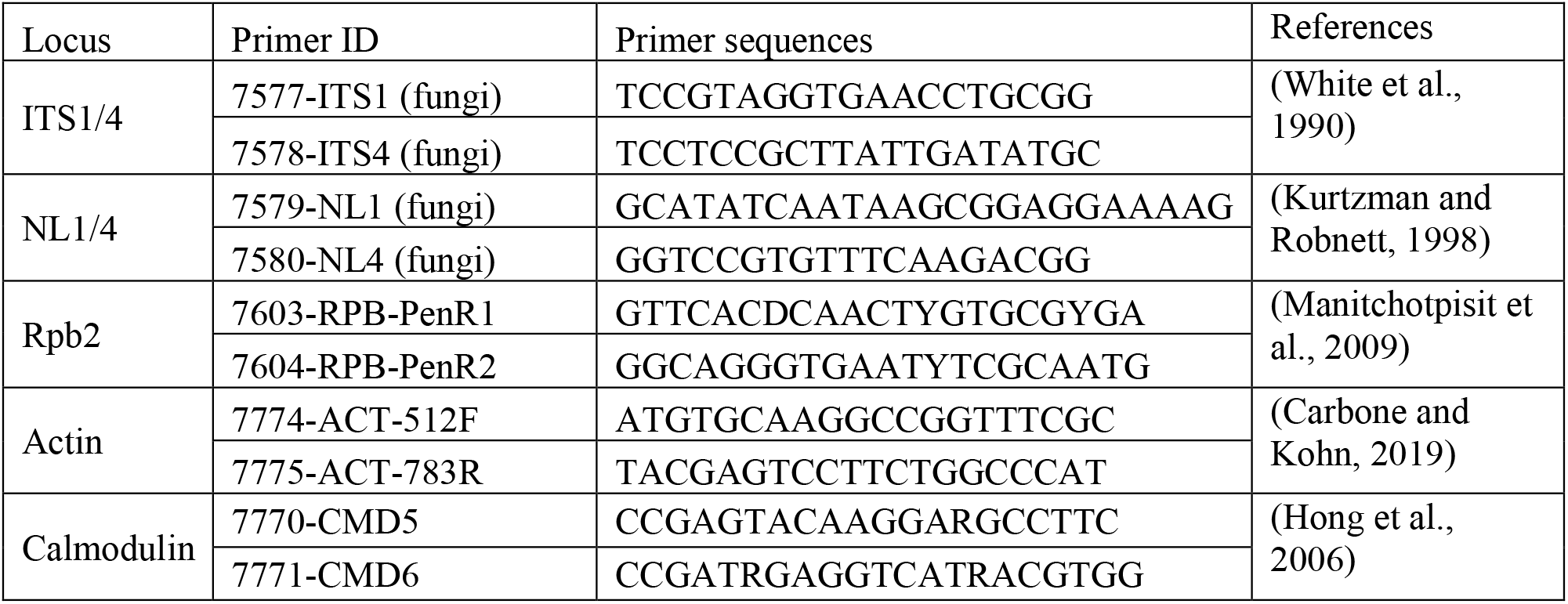
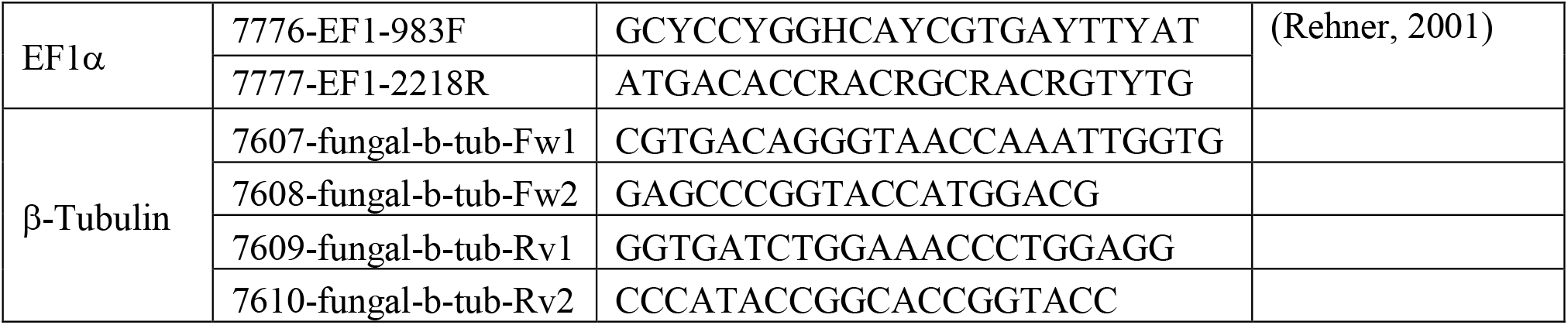
Primers used in this study.

**Table 2.**
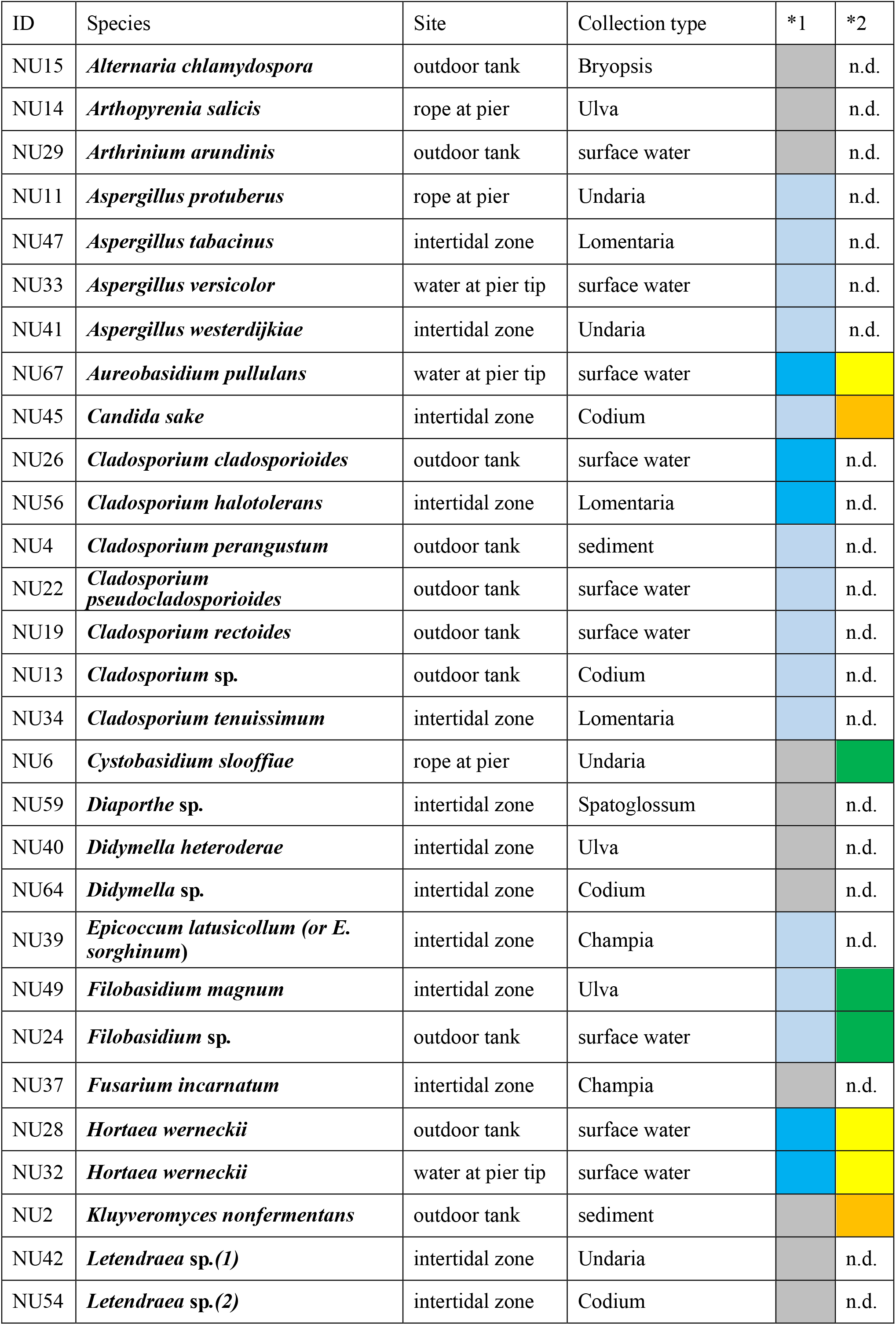
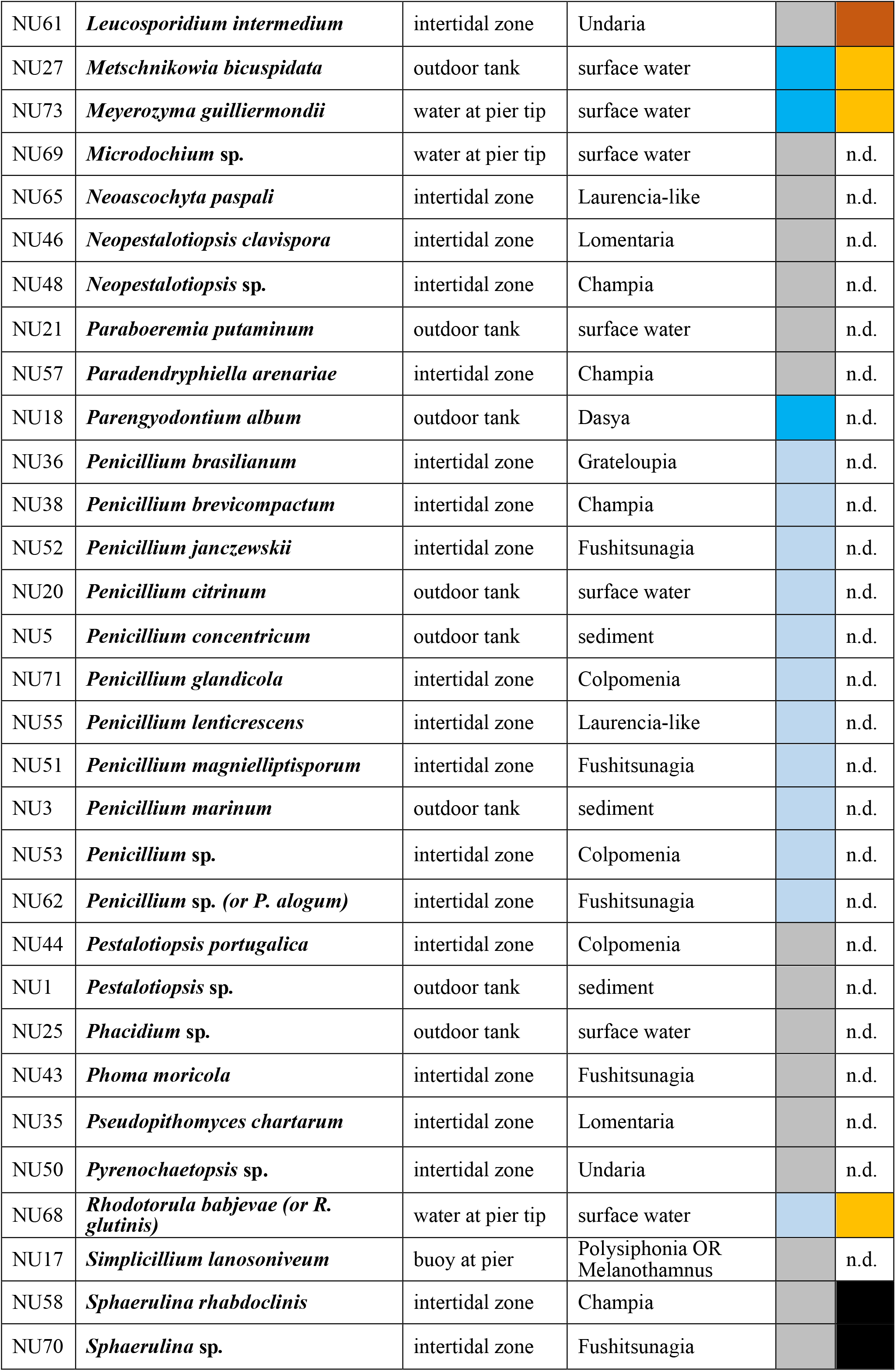
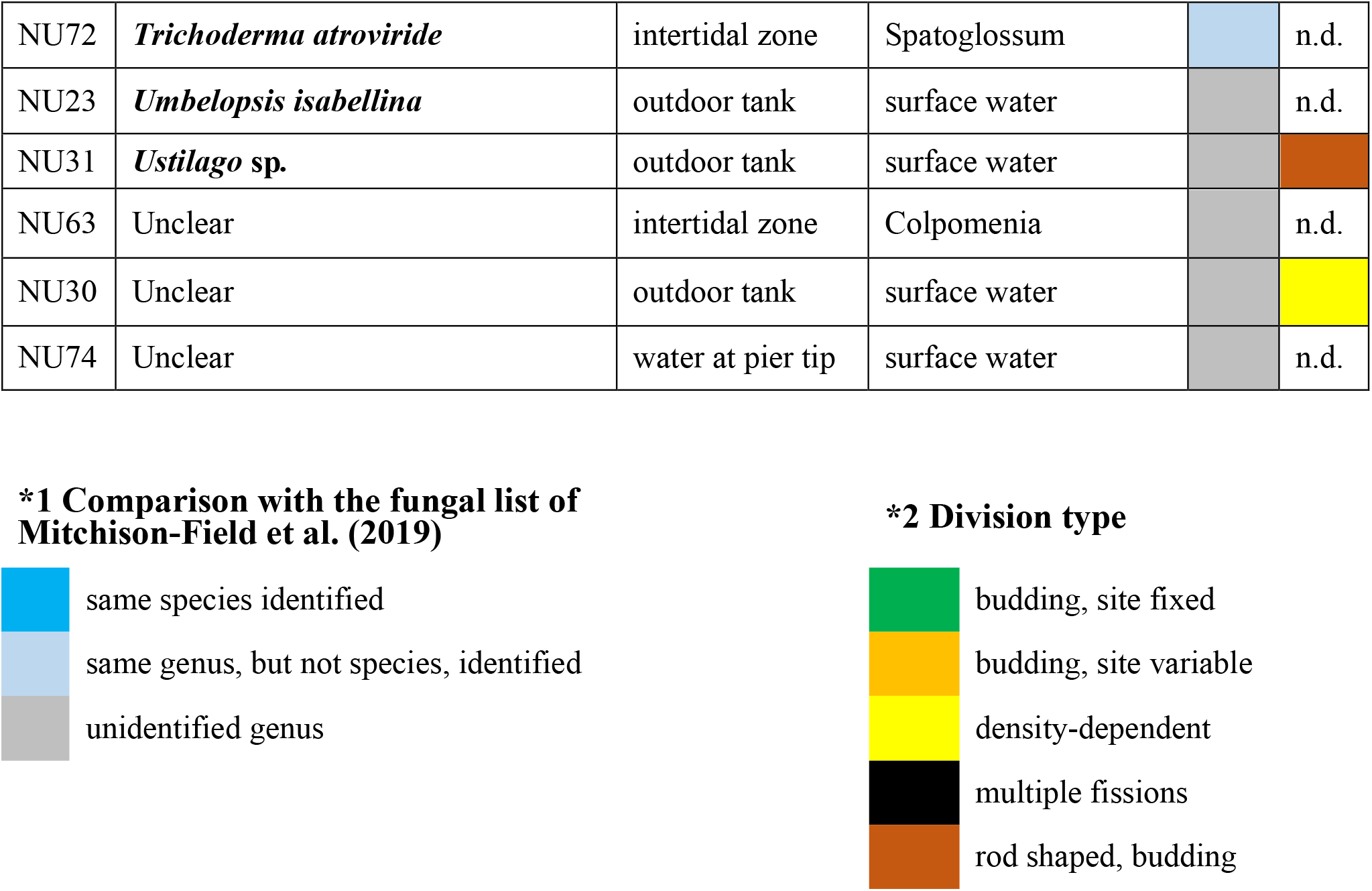
List of marine fungi identified in this study (alphabetical order)

### Comparison with the Woods Hole collection

There were considerable differences between the current collection at Sugashima and what was identified around Woods Hole (Mitchison-Field et al., 2019) (Table 2, *1). Only seven species were isolated in both studies (*Aureobasidium pullulans, Cladosporium cladosporioides, Cladosporium halotolerans, Hortaea werneckii, Metschnikowia bicuspidate, Meyerozyma guilliermondii*, and *Parengyodontium album*). At the genus level, only 13 genera were common (*Aspergillus, Candida, Epicoccum, Filobasidium, Penicillium, Rhodotorula*, and *Trichoderma*, in addition to aforementioned six genera), while other genera were uniquely isolated in either location: 27 from Sugashima (*Alternaria, Arthopyrenia, Arthrinium, Cystobasidium, Diaporthe, Didymella, Fusarium, Kluyveromyces, Letendraea, Leucosporidium, Microdochium, Neoascochyta, Neopestalotiopsis, Paraboeremia, Paradendryphiella, Pestalotiopsis, Phacidium, Phoma, Pyrenochaetopsis, Simplicillium, Sphaerulina, Umbelopsis, Ustilago*, and four unclear genera), and six from Woods Hole (*Apiotrichum, Cadophora, Cryptococcus, Exophiala, Knufia, Phaeotheca*). While this discrepancy may partly stem from seasonal or regional differences, it more likely reflects the limited coverage of fungi in both studies. The current survey agrees with the view that many more species exist in the ocean (Amend et al., 2019).

### Budding yeast species with fixed and variable bud positions

Fifteen fungal species produced yeast-like colonies on culture plates. Live imaging was performed for these species at least thrice at different cell densities. Five of them turned out to have *S. cerevisiae*-like budding-growth cycles, where a daughter cell emerged from the round-shaped mother cell and a mother cell can produce a second daughter at other sites than the previous one (*Candida sake, Kluyveromyces nonfermentans, Metschnikowia bicuspidate, Meyerozyma guilliermondii, Rhodotorula* sp.) (Fig. 3A, Video 1). Interestingly, the budding sites of three other yeasts were fixed, although the mother cell had a nearly round shape (*Cystobasidium slooffiae, Filobasidium magnum, Filobasidium* sp.) (Fig. 3B, Video 1). Four other fungi were rod-shaped, in which budding and/or filamentous growth were observed (*Leucosporidium intermedium, Ustilago* sp., *Sphaerulina rhabdoclinis, Sphaerulina* sp.) (Video 2).

**Figure 3.**
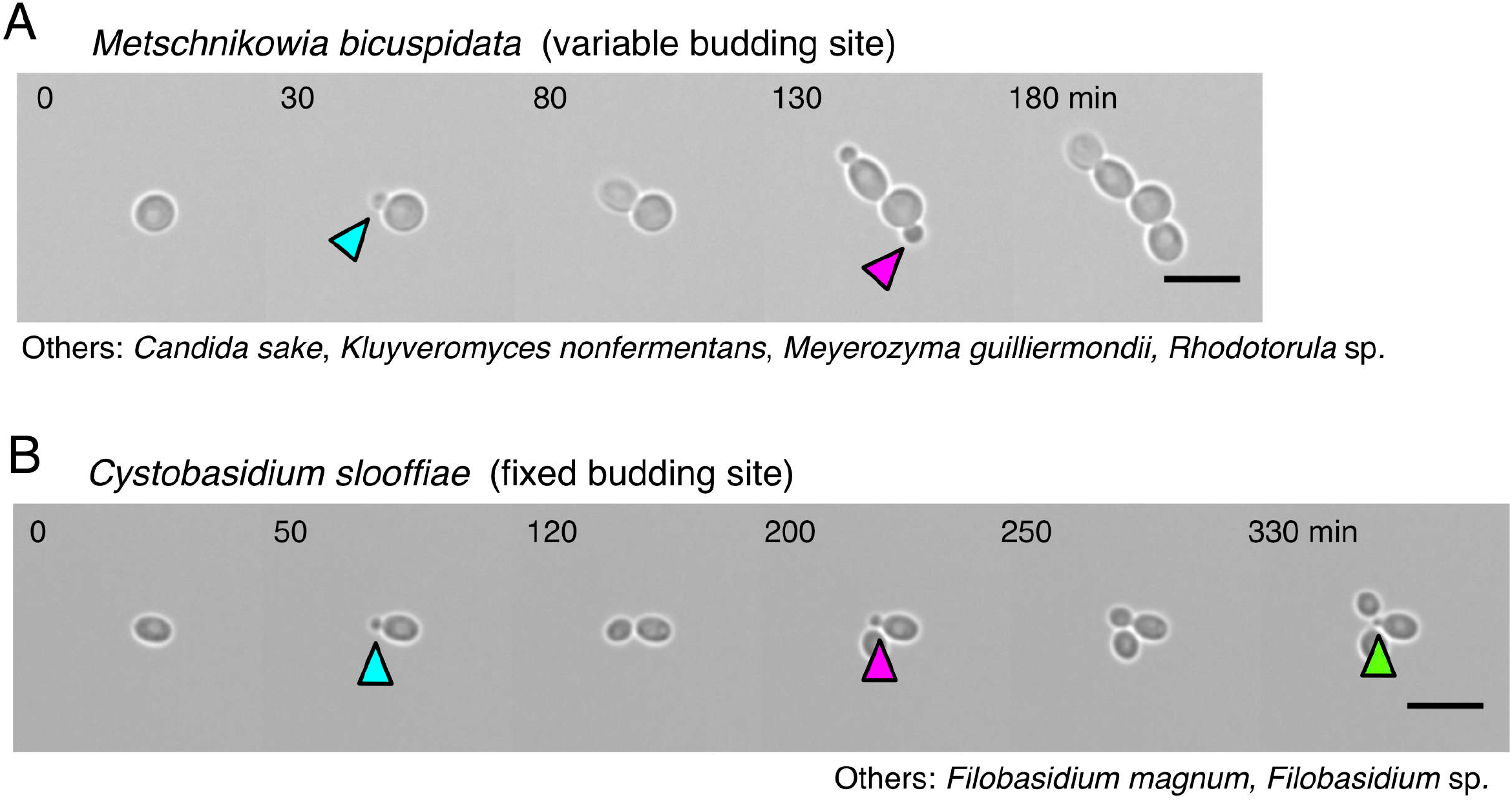
Budding yeast has either a fixed or variable budding site. (A) *S. cerevisiae*-type budding yeast, where a bud emerges at seemingly random sites on the round mother cell. (B) The budding site is fixed at one site of the mother cell. Blue arrows, initially emerged bud; magenta arrow, second bud; green arrow, third bud. Scale bars, 10 µm.

### Cell density-dependent division pattern alteration in the black yeast *A. pullulans*

The remaining three fungi that formed yeast-like colonies produced black or brown pigment. Two of them have been extensively studied by (Mitchison-Field et al., 2019): *A. pullulans* (Fig. 4A) and *H. werneckii* (Fig. 5A). *A. pullulans* in (Mitchison-Field et al., 2019) had a round shape and showed multiple buds at one site. However, in my first few imaging attempts, such a characteristic division pattern was never observed. Instead, they first grew as filaments, and later produced multiple buds from many locations on the filament. In the course of repetitive imaging, it was noticed that a division pattern closer to that described in (Mitchison-Field et al., 2019) could be obtained when the initial cell density was increased. To test the possibility that the division pattern is density-dependent, a 10-fold dilution series of the strain was prepared for imaging (relative cell densities: ×1, ×10, ×100, and ×1000). When the initial cell density was low, a single cell first elongated with occasional septation, followed by multiple budding from the elongated cell filament (100%, n = 65) (Fig. 4B, D, Video 3). The higher the cell density, the faster the bud emerged (Fig. 4C). Immediate budding without extensive mother cell elongation or septation was observed only when the initial cell density was high (Fig. 4B, 19 out of 30 cells). However, the majority of the cells (14/19) showed buds at both poles of the mother cell (Fig. 4E) rather than produced buds from the same pole, as reported in (Mitchison-Field et al., 2019) (5/19) (Fig. 4F). Thus, this black yeast species showed division pattern variation depending on the cell density.

**Figure 4.**
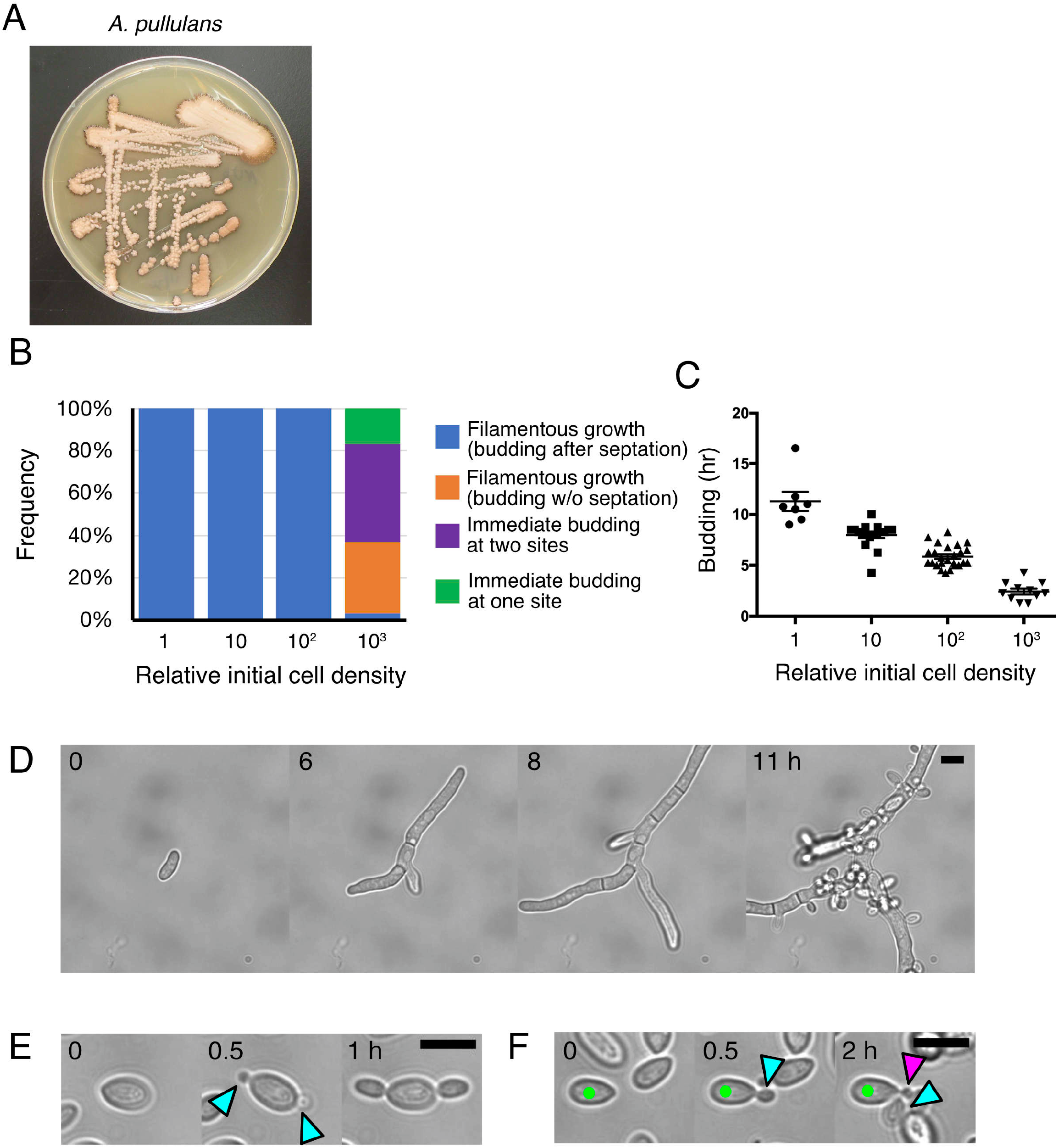
Division pattern variation in the black yeast *Aureobasidium pullulans*. (A) *A. pullulans* colonies. The peripheries of the colonies turned brown after prolonged storage at 4 °C (fresh colonies are uncoloured). (B) Four distinct bud formation patterns dependent on the initial cell density. Saturated cultures were diluted at four different concentrations, and the growth and budding style was assessed (n = 22, 18, 25, 30 [left to right]). (C) Timing of bud emergence after a cell started to grow in a filamentous manner (± SEM) (n = 7, 18, 25, 11 [left to right]). The one-way ANOVA detected significant differences between groups (F = 77.06, p < 0.0001). The post-hoc test was performed by Tukey’s test (p < 0.0001 for each comparison). (D) Filamentous growth with occasional septation and branching, followed by budding from various sites on the filament. This mode of growth/budding was dominant when the initial cell density was low. (E, F) Immediate budding without apparent cell growth or septation. Two buds simultaneously emerged at two opposite sites in (E), whereas two buds sequentially emerged at one site of the mother cell in (F). Blue arrows, initially emerged buds; magenta arrow, second bud; green, mother cell. Scale bars, 10 µm.

An earlier study involving immunofluorescence microscopy of microtubules and actin of other *A. pullulans* strains also showed filamentous morphology (Kopecka et al., 2003). Therefore, time-lapse imaging of *A. pullulans* strain collected in (Mitchison-Field et al., 2019) was performed in an identical condition to that for the Sugashima strain. Interestingly, the Woods Hole strain also showed filamentous growth at low density (100%, n = 30) and immediate multi-budding was observed only at the highest density. These results suggest that density-dependency of growth/division mode alteration is a common feature of *A. pullulans*.

### Cell density-dependent division pattern alteration in the black yeast *H. werneckii*

Two clones of *H. werneckii* were isolated, and their colony sizes were slightly different (Fig. 5A). The barcode sequences had high similarity but were not identical (1 bp mismatch). These were interpreted as natural variants of the same species. Imaging of these clones initially produced puzzling results: the reported characteristic division pattern– a single cell almost always undergoes fission, followed by budding (Mitchison-Field et al., 2019) – was rarely observed. To test the possibility that, similar to *A. basidium*, the initial cell density might have affected the division pattern, the number of inoculated cells was varied (×1, ×10, ×100, ×300). The initial cell numbers dramatically affected the mode of the first few cell divisions in both isolates (Fig. 5B presents the result of NU28; Video 4). Multiple rounds of septation were observed when cell density was low and multiple budding occurred hours later (Fig. 5B, D). The higher the cell density, the earlier the bud emerged (Fig. 5C). In contrast, the reported mode of division – fission followed by budding – was predominantly observed when the cell density was high (Fig. 5B, E). However, cells that formed buds without fission were also observed, which was not reported in (Mitchison-Field et al., 2019) (Fig. 5B, F). The strain collected in (Mitchison-Field et al., 2019) also showed filamentous growth at the lowest density in the liquid culture (100%, n = 35). Thus, the division pattern variability of *H. werneckii* was similar to that of *A. pullulans*. The finding corroborates the observation of the colonies on the plate for >30 *H. werneckii* strains, where both filamentous and yeast-like cells are identified (Zalar et al., 2019).

**Figure 5.**
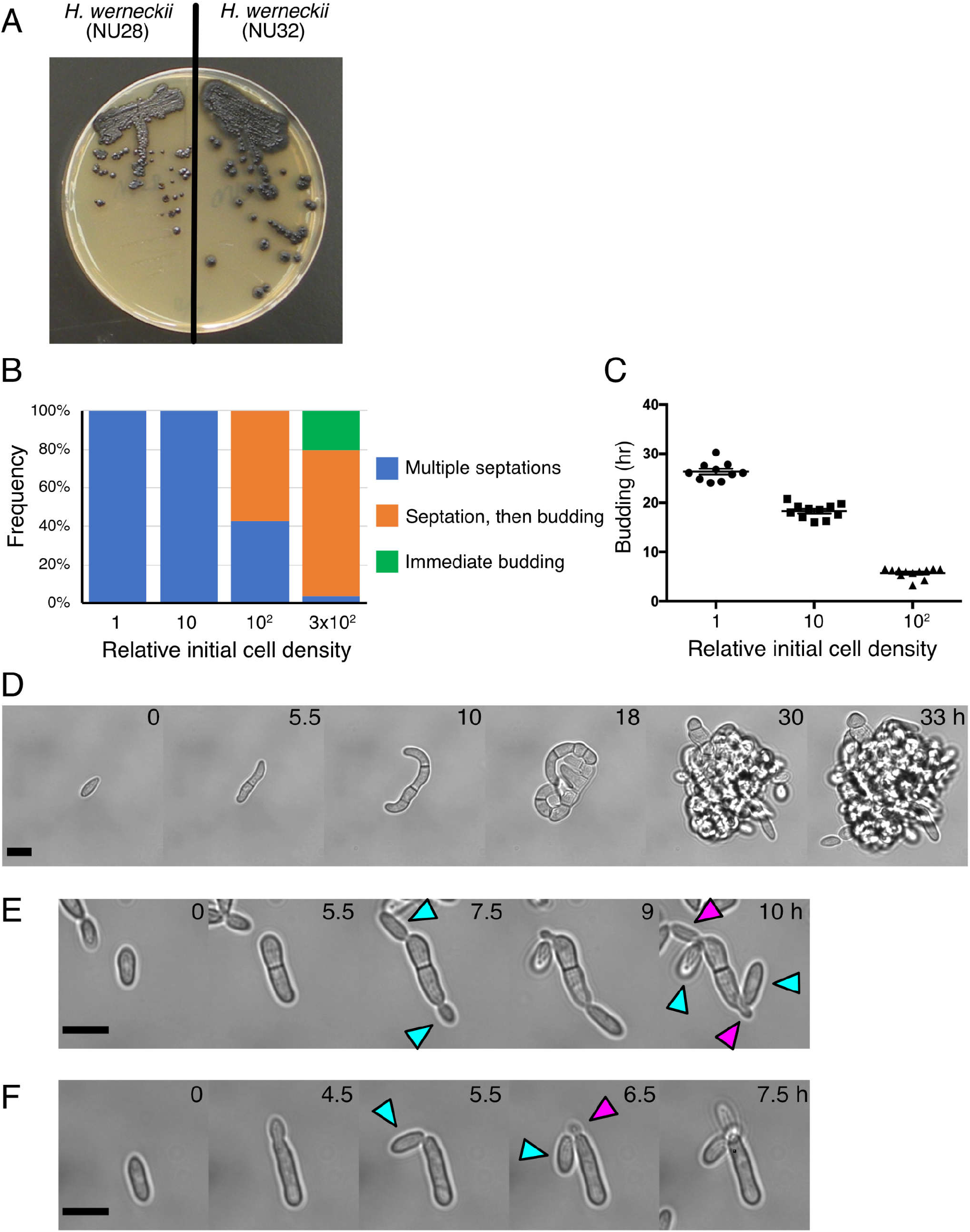
Division pattern variation in the black yeast *Hortaea werneckii*. (A) Colonies of two *H. werneckii* strains, which had different growth rates. (B) Three distinct bud formation patterns dependent on the initial cell density. Saturated cultures were diluted at four different concentrations, and the growth and budding style was assessed (n = 30, 11, 28, 54 [left to right]). (C) Timing of bud formation after the first septation (± SEM) (n = 19, 11, 12 [left to right]). The one-way ANOVA detected significant differences between three groups (F = 549.2, p < 0.0001). The post-hoc test was performed by Tukey’s test (p < 0.0001 for each comparison). (D) Filamentous growth with septation and branching, followed by budding from various sites on the curved filament (a released bud is seen at lower-left corner at 43 h). This mode of growth/budding was dominant when the initial cell density was low. (E) Elongation, septation, followed by budding. Multiple buds sequentially emerged from a “mother” cell that had undergone septation. This mode of septation/budding was dominant when the initial cell density was high. (F) Immediate budding without septation. Multiple buds sequentially emerged from a “mother” cell. Blue and magenta arrows indicate the first and second buds, respectively. Scale bars, 10 µm.

### Cell density-dependent division pattern alteration in an unidentified black yeast species

In the present study, another yeast strain (NU30) formed dark brown-coloured colonies on the plate (Fig. 6A). The barcode sequences did not perfectly match any registered species; because the mismatch was large, the name of this yeast could not be determined. To reveal its growth/division pattern and test if it was altered depending on the initial cell density, time-lapse imaging was conducted at four different initial cell densities (×1, ×10, ×100, and ×700). The cell division pattern observed was remarkably similar to that of *H. werneckii* (Fig. 6B–F, Video 5). Budding occurred after multiple septations when the initial cell density was low, whereas a mother cell, without cell septation, produced multiple buds sequentially from the same site (blue and magenta arrows) when the initial cell density was high (Fig. 6E, F).

**Figure 6.**
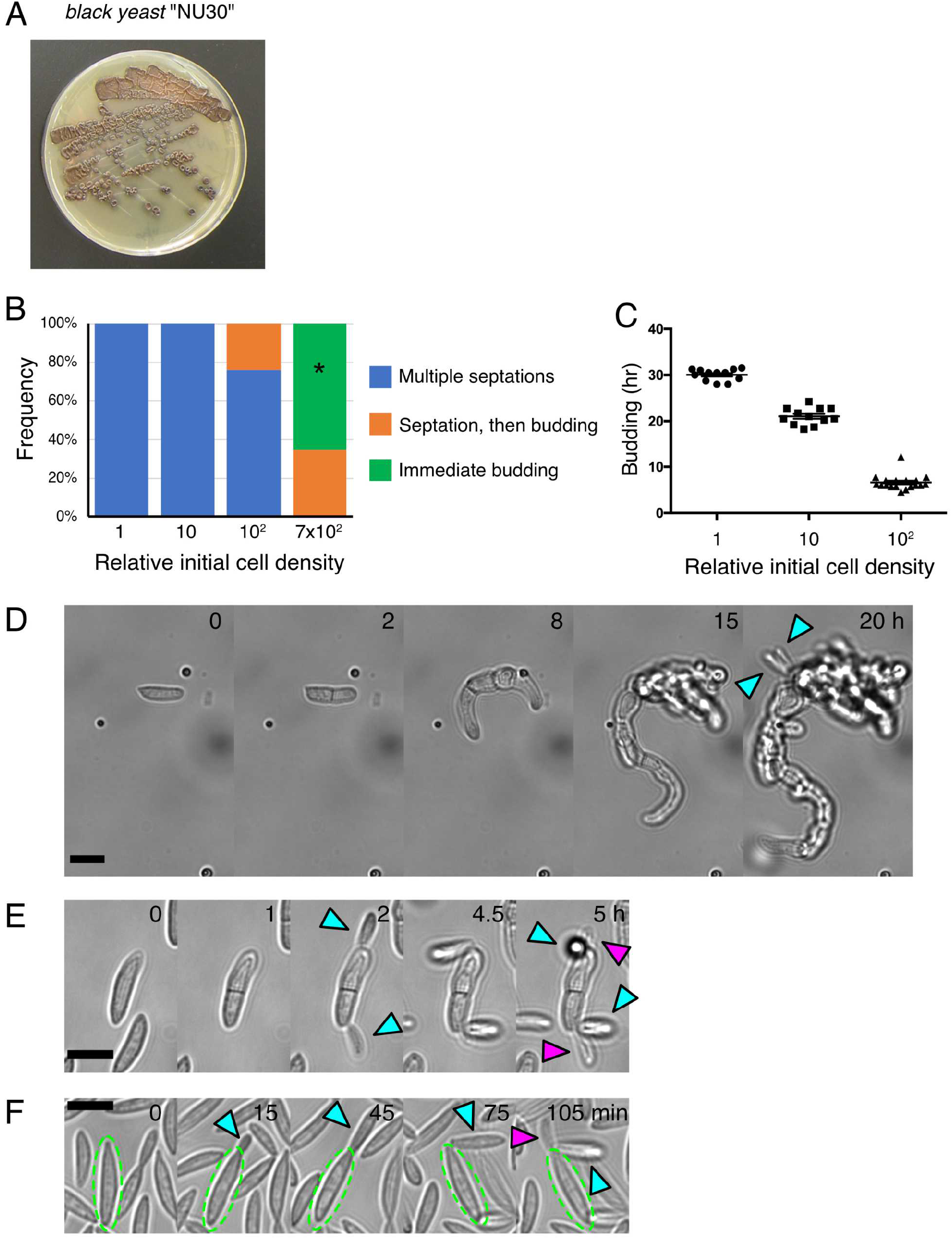
Division pattern variation in an unidentified black yeast species “NU30”. (A) Colonies of NU30. Each colony turned dark brown after prolonged storage at 4 °C (fresh colonies were uncoloured). (B) Three distinct bud formation patterns dependent on the initial cell density. Saturated cultures were diluted at four different concentrations, and the growth and budding style was assessed (n = 25, 12, 25, 37 [left to right]). Asterisk: the frequency of immediately budding cells was underestimated, as many cells did not stick to the glass and were hard to count, whereas all the cells that underwent septations were immobile and countable. (C) Timing of bud formation after the first septation (± SEM) for the cells that formed multiple septations (n = 12, 12, 19 [left to right]). The one-way ANOVA detected significant differences between groups (F = 811.7, p < 0.0001). The post-hoc test was performed by Tukey’s test (p < 0.0001 for each comparison). (D) Filamentous growth with septation and branching, followed by budding from various sites on the curved filament (20 h). This mode of growth/budding was dominant when the initial cell density was low. (E) Single septation, followed by budding. Multiple buds sequentially emerged from a “mother” cell that had undergone septation. (F) Immediate budding without septation. Multiple buds sequentially emerged from the “mother” cell (green). This mode of budding was dominant when the initial cell density was high. Blue and magenta arrows indicate the first and second buds, respectively. Scale bars, 10 µm.

### Nuclear segregation scales with cell length in *H. werneckii*

In the model budding yeast *S. cerevisiae*, budding starts at the G1/S transition of the cell cycle, and nuclear division takes place when the bud reaches a certain size (Juanes and Piatti, 2016). In *S. pombe*, the nucleus is kept in the centre of the cell during interphase (mostly G2 phase), then mitotic commitment occurs when the cell reaches a certain length (∼14 µm) (Wood and Nurse, 2015). In both cases, cell division (septation) occurs immediately after nuclear division, producing daughter cells that have an identical shape to the mother. In contrast, in *A. nidulans* and *Ashbya gossypii*, the model filamentous fungi, multiple nuclear divisions take place without septation in the tip-growing cells, resulting in the production of multinucleated cells (Fischer, 1999; Gladfelter et al., 2006). It was curious whether nuclear and cell divisions were coupled in the above three black yeast species, whose division pattern was flexible.

To follow nuclear division and septation/budding in live cells, their nuclei were stained with Hoechst 33342, which is known to be permeable in many cell types, including *S. pombe* (Haraguchi et al., 1999). However, staining and live imaging of *A. pullulans* and NU30 were not successful. In contrast, *H. werneckii* was stainable, and its nuclear dynamics was observable in real time (Fig. 7, Video 6, 7). Therefore, time-lapse imaging with Hoechst 33342 using a spinning-disc confocal microscope was performed for *H. werneckii*.

**Figure 7.**
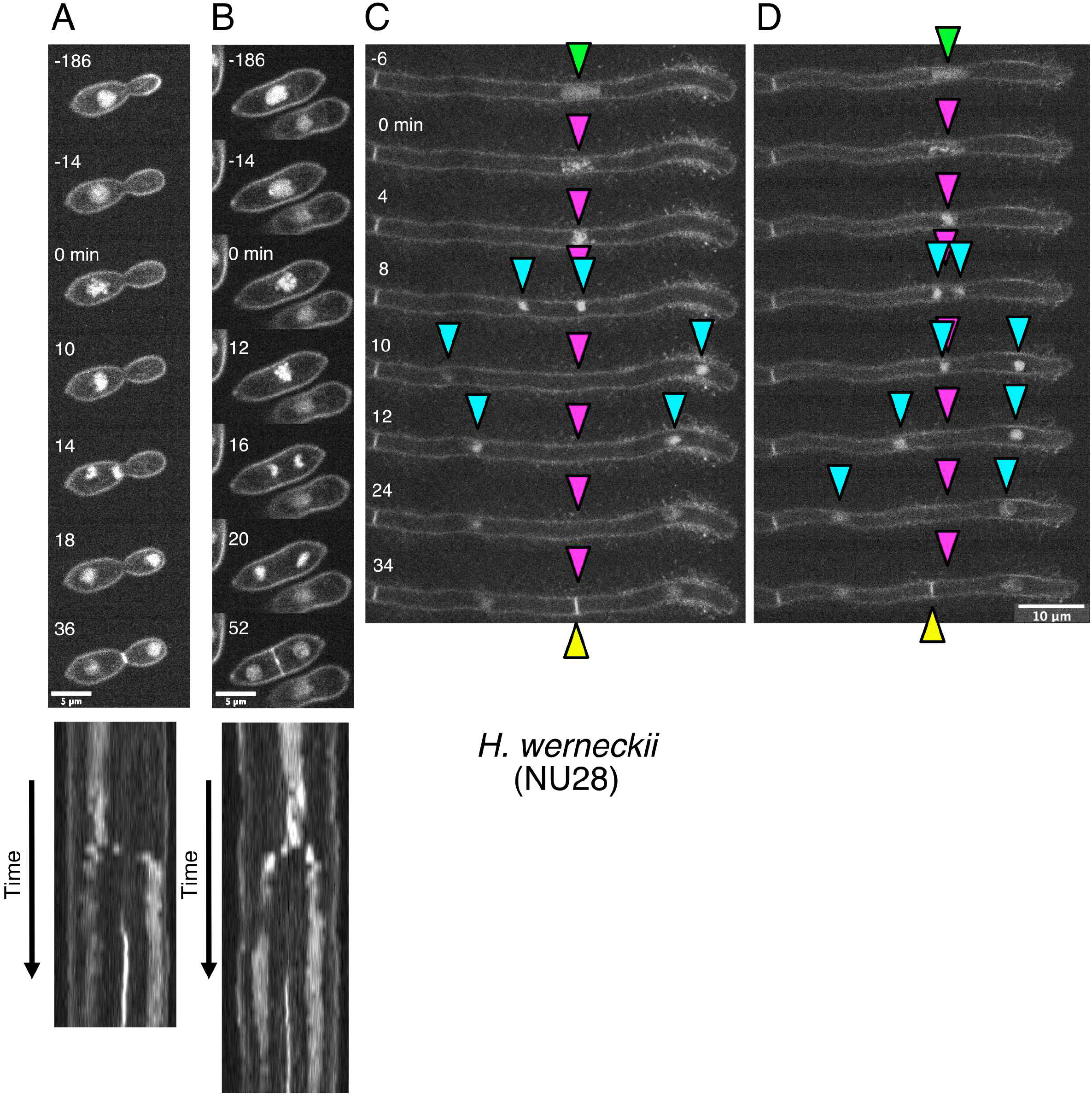
Nuclear dynamics in *H. werneckii*. (A) Nuclear dynamics during budding-type cell division. The corresponding kymograph is shown at the bottom. Time 0 indicates mitotic entry. (B) Nuclear dynamics during fission-type cell division. The corresponding kymograph is shown at the bottom. Time 0 indicates mitotic entry. (C, D) Nuclear dynamics in tip-growing cells. Green arrowheads, interphase nuclei; magenta, position of condensed chromosomes at the mitotic entry; blue, sister chromatids/nuclei; yellow, septum. Septum was formed at the site of mitotic chromosomes in (C), whereas it was deviated in (D). Segregation of sister chromatids is asymmetric relative to the metaphase chromosome.

In the budding type of mitosis shown in Fig. 7A, which prevailed when the cell density was high, nuclear separation occurred when the bud size reached 60 ± 11 % (± SD) of the mother (n = 30). The nucleus was positioned in the mother, slightly on the bud neck side at the time of chromosome condensation (i.e. mitotic entry) (44 ± 6 % position from the bud neck). Sister chromatid separation occurred in the mother, and one of the sister chromatids moved into the bud. Kymograph analysis indicated that the sister chromatid motility relative to the cell edge was asymmetric (Fig. 7A, bottom): the sister on the bud side moved much longer distances, whereas the sister in the mother moved much less or sometimes showed bud-oriented movement. The maximum distance between sister chromatids was 9.4 ± 1.1 µm, which was comparable to the mother cell length (9.2 ± 1.0 µm) and was 36 % shorter than the entire cell length (i.e., daughter + mother length). These results suggested that the anaphase spindle was motile in the cell. The mitotic duration (chromosome condensation to anaphase onset) was 14 ± 7 min. Septation, which was indicated by a straight line of the Hoechst dye at the bud neck, was completed 16 ± 6 min after sister chromatid separation.

In the fission type of mitosis shown in Fig. 7B, the nucleus was positioned near the centre of the cell at the time of chromosome condensation (16 or 24 out of 26 cells had the nucleus at 45–55% or 40–60% position along the cell axis, respectively). After 16 ± 3 min (n = 26), the sister chromatids were separated. Unlike in *S. pombe*, chromosome segregation did not always occur symmetrically (232–236 min time point in Video 6) and the chromatids rarely reached the cell edge (maximum sister chromatid distance was 5.6 ± 0.5 µm, which was 54 ± 4 % of cell length) (Fig. 7B, bottom). Septation was completed in the middle of the cell at 28 ± 4 min after sister chromatid separation.

In the tip-growing cells, which were observed at low cell density, a single nucleus was detected, and it moved apically during tip growth, contrary to the multiple nuclei in *A. nidulans* or *A. gossypii* (Fig. 7C and 7D; flare-like structures were also stained on the cell surface by Hoechst 33342). Therefore, the nucleus stayed near the centre of the cell (47 ± 5 % position from the tip at mitotic entry, n = 32). The cell length was quite variable at the time of chromosome condensation (36 ± 11 µm, ±SD, n = 32); in some cases, it reached >40 µm, which was four times longer than the rod-shaped or budding cells described above. Mitotic duration was 8.1 ± 1.2 min (n = 27). Sister chromatids moved longer distances (53 ± 6 % of the cell length, n = 31) and septation occurred in the middle of the two nuclei at 21 ± 3 min (n = 26) after sister chromatid separation. However, the middle part was motile during anaphase, implying that the anaphase spindle was motile in the cell. Consequently, the chromosome position at metaphase was not always consistent with the septation position (Fig. 7D).

Thus, despite variations in cell size and geometry (rod, filament, bud), septation was coupled with nuclear division. However, the segregating distance of sister chromatids varied significantly, scaling with the cell length and shape. This suggests that the mechanics of the cell division apparatus are adjusted to each mode of cell growth.

## Discussion

Two major conclusions can be drawn from this study. First, a very limited local survey has increased the list of marine-derived fungi, illustrating their diversity in the ocean. The species could not be determined for several fungi, suggesting that they might represent undescribed species. More comprehensive sampling and taxonomic analyses are needed to further expand the list of fungi from the ocean.

Second, all three collected black yeast species changed their division mode depending on the cell density. This plasticity may be beneficial for them, particularly when they reside on the surfaces of animals and macroalgae. Filamentous growth with branching is an excellent means to explore unoccupied areas (Coudert et al., 2019), whereas budding in a crowded environment allows the clone to be released to the free water and translocated to other animal/algae surfaces. This property somewhat resembles that of filamentous fungi such as *Aspergillus*; they exhibited 2D hyphal growth, followed by conidia (asexual spores) release (Adams et al., 1998). Switching between yeast-like budding and hyphal growth is also observed for pathogenic, dimorphic fungi such as *Candida albicans* (Merson-Davies and Odds, 1989; Sudbery et al., 2004; Sudbery, 2011). However, the filament was curved and area exploration was not optimised for *H. werneckii* and NU30; hence, the advantage of this growth/division mode remains unclear.

Combined with this and previous studies using live-cell microscopy (Mitchison-Field et al., 2019), all five observed black yeast species showed at least two growth/division patterns. The cell density-dependent plasticity might lie in nutritional states. For example, some *H. werneckii* strains show different morphology on the plate depending on the presence or absence of NaCl in the medium (Zalar et al., 2019). Alternatively, a quorum-sensing mechanism (Albuquerque and Casadevall, 2012) might be responsible for the inhibition of filament formation at high cell densities. Elucidating the chemical and/or physical cues that trigger the division mode alteration and the prevalence of plasticity in marine fungi would be interesting topics for future research.

## Materials and methods

### Isolation of marine fungi at NU-MBL

Samples were collected for 8 days from 4^th^ March to 5^th^ June 2020, in which the majority were obtained on the 22^nd^ and 23^rd^ of April. The ocean water temperature was 15.5 °C and the salinity was 34.9 ‰. This salinity indicates that there was no significant fresh water flow into this area from the large rivers in Ise Bay. Surface water samples were collected at the pier of the NU-MBL using a plastic bucket (Lat. 34°29′8″N, Long. 136°52′32″E) (Fig. 1B, green). One litre of seawater was filtered using a 0.45-µm Millipore Stericup to obtain a 100× concentration. One millilitre of the concentrate was spread onto three types of media, which were similar to those used in (Mitchison-Field et al., 2019): YPD (10 g/L yeast extract, 10 g/L peptone [Bacto tryptone], 20 g/L glucose, 12 g/L agar in seawater), malt medium (20 g/L malt extract, 6 g/L tryptone, 20 g/L glucose, 12 g/L agar in seawater), and potato dextrose (24 g/L potato dextrose broth, 12 g/L agar in seawater). All species tested could grow on any medium; retrospectively, the preparation of three different media was not needed. Sediment samples were collected once by taking the mud at the bottom of the outside tank (Fig. 1C). Seaweeds were collected from the intertidal zone in front of the NU-MBL (Fig. 1B, magenta). Approximately 3 cm × 3 cm fragments of seaweed were obtained, which were rinsed with semi-sterile seawater (∼500 mL) three times, followed by knife-cutting into small pieces and spreading onto a plate (Fig. 1D). All media were made with seawater at NU-MBL, which was pre-filtered with ADVANTEC filter paper #2. Antibiotics were added to the medium to avoid bacterial growth (20 µg/mL carbenicillin, 100 µg/mL chloramphenicol, 10 µg/mL tetracycline) (Mitchison-Field et al., 2019). All fungal isolates were stored at - 80 °C in YPD medium containing 20% glycerol.

### PCR and sequencing

The ITS and NL sequences were sequenced for all strains using the primers listed in Table 1. If the species could not be determined through these two sequences, other loci were amplified and sequenced (Rpb2, actin, calmodulin, EF1α, β-tubulin). PCR was performed with the KOD-ONE kit (TOYOBO) using a colony or the extracted genomic DNA as the template. DNA extraction was performed by boiling a piece of fungal colony for 5 min in 0.25% SDS followed by table-top centrifugation.

### Species identification

Sequence homology was determined using the BLASTN program. If both ITS and NL sequences were perfectly matched to a single species, the fungal isolate was concluded to be the species. In some cases, either the ITS or NL sequences had a 100% match to a certain species, whereas the other did not show an exact match. When the mismatch was less than 3 bp, the fungal isolate was likely the species with a slight strain-specific sequence variation, whereas a >3 bp mismatch in two or three barcodes led to the assignment of possible novel species or genera. If a species could not be specified with ITS and NL, the sequences of β-tubulin, Rpd2, actin, calmodulin, and EF1α were also used, dependent on the genus. For example, when ITS and NLS sequences perfectly matched several *Cladosporium* species, the actin locus was further sequences and specified the species. Calmodulin sequences were used for *Penicillium*. All the barcode sequences are presented in Table S1.

### Live microscopy

Initially, cells grown on the YPD plate were directly inoculated into YPD liquid medium in an 8-well glass-bottom dish (Iwaki). However, the division modes were not consistent between experiments for some species. Therefore, cells were grown in a YPD liquid medium until saturation and inoculated into an 8-well plate after cell counting (300 µL culture volume). Note that this liquid-based imaging condition is different from that in (Mitchison-Field et al., 2019), in which agar pads were used. The growths of 10^3^, 10^4^, 10^5^, and >10^5^ cells were compared for black yeasts. Transmission light imaging was carried out at 23 °C with a Nikon inverted microscope (Ti) with a 40× 0.95 NA lens (Plan Apo) and a Zyla CMOS camera (Andor). The brightness and contrast of the obtained images were adjusted using FIJI software. The statistical analyses were performed using the GraphPad Prism software. For DNA imaging of *Hortaea werneckii*, cells were cultured with a more transparent medium. Synthetic minimal medium, similar to what was used for *Aspergillus nidulans*, was made with seawater (6 g/L NaNO3, 0.52 g/L KCl, 1.52 g/L KH2PO4, 10 g/L glucose, 1.5 mL trace elements, 10 mg/L biotin, 0.25 g/L MgSO4, pH = 6.5 [adjusted with NaOH]) (Edzuka et al., 2014). Hoechst 33342 was added at 3–6 µg/mL (final). DNA imaging was performed with another Nikon inverted microscope (Ti2) attached to a 100× 1.40 NA lens, a 405-nm laser, CSU-10 spinning-disc confocal unit, and the CMOS camera Zyla (at 23 °C). The time intervals used were mostly 2 min, but sometimes 2.5 min or 5 min was used. The microscopes were controlled using NIS Elements software (Nikon).

## Acknowledgements

I am grateful to Tomoya Edzuka and Maki Shirae-Kurabayashi for seaweed collection, Masahiro Suzuki (Kobe University) for help with seaweed identification, Masashi Fukuoka for daily temperature and salinity measurements, Naoto Jimi for helpful comments on taxonomy, Amy Gladfelter (Marine Biological Laboratory, Woods Hole/University of North Carolina) and Christine Field (Marine Biological Laboratory, Woods Hole/Harvard Medical School) for yeast strains, and Amy Gladfelter for the introduction to marine fungal biology and encouraging sample collection on the Japanese coast. This work was supported by JSPS KAKENHI (17H06471, 18KK0202).

## Supplementary video legends

**Video 1. Cell division of marine-derived budding yeast**

Time-lapse imaging of two budding yeast species (5-min intervals). The budding site is variable (left) or fixed (right) depending on the species.

**Video 2. Cell division of rod-shaped fungi *Ustilago* sp**., ***Leucosporidium intermedium*, and *Sphaerulina rhabdoclinis***

Time-lapse imaging of three fungi that have a rod shape and form yeast-like colonies on the plate (5-min interval).

**Video 3. Cell division of the black yeast *Aureobasidium pullulans***

Live imaging of *Aureobasidium pullulans* (NU67) at four different initial cell densities. Images were acquired every 15 min.

**Video 4. Cell division of the black yeast *Hortaea werneckii***

Live imaging of *Hortaea werneckii* (NU28) at three different initial cell densities (lower left, high: upper left, medium: right, low). Images were acquired every 15 min.

**Video 5. Cell division of an unidentified black yeast species**

Live imaging of NU30 (unnamed) at three different initial cell densities (lower left, high: upper left, medium: right, low). Images were acquired every 15 min.

**Video 6. Nuclear dynamics in the black yeast *Hortaea werneckii – budding and fission*** Live imaging of chromosomes in *Hortaea werneckii* (NU28) during budding-and fission-type mitosis. Images were acquired every 2 min.

**Video 7. Nuclear dynamics in the black yeast *Hortaea werneckii – growing tip***

Live imaging of chromosomes in *Hortaea werneckii* (NU28) during mitosis of tip-growing cells. Images were acquired every 5 min. Note that flare-like structures are also stained.

